# The double-edged sword of urban life: Invasive grey squirrels perceive foraging as safer close to roads under noisier conditions, yet riskier when noise is inconsistent

**DOI:** 10.1101/2024.05.31.596847

**Authors:** Kristin P. J. Thompson, Sasha R.X. Dall

## Abstract

1. Mortality risk influences decisions of foraging animals such that foraging and feeding behaviours are usually biased towards habitats where benefits of feeding outweigh risks. Human activity and associated disturbance are thought to be perceived as a source of risk analogous to predation risk in wildlife. As such, can alter behaviour and habitat use of foraging animals.
2. Urban wildlife faces increased exposure to human disturbance, meaning that they may face an increase in perceived risk during food acquisition. Urban habitats also include novel and altered food resources that could result in urban wildlife having to face distinct trade-offs associated with foraging and patch use compared to rural counterparts.
3. To examine how a relatively successful invasive urban mammal, the eastern grey squirrel, balances risk and safety under human disturbance, we measured giving-up densities (GUD) at artificial food patches placed at sites subject to varying levels of urbanisation to investigate how features associated with human disturbance might influence feeding decisions.
4. We found that differences in GUDs between ‘safe’ and risky’ patches were reduced closer to roads only under noisy conditions, suggesting that the risk of predation is perceived by squirrels as reduced when disturbance from human activities is highest. However, there was also a significant effect of the variability in noise levels on patterns of patch exploitation, with higher GUDs and larger GUD differences between safe and risky patches found under more variable noise levels, suggesting that squirrels might also find dynamic human disturbances startling or distracting while foraging.
5. Synthesis: Our results show how human activities can have doubled-edged impacts on the urban landscape of fear through offering reduced risk from predators whilst increasing foraging costs via noise disturbance. Future research could consider how foraging decisions in urban wildlife vary according to food supply and disturbance levels, to shed further light on foraging trade-offs made by wildlife under increasing urbanisation levels.

## Introduction

Balancing costs and benefits associated with food acquisition is key for survival for foraging animals, therefore, decisions of where, when, and how long to forage are shaped by the animal’s assessment of risk at a patch (Brown, 1999; Brown, 1988). Feeding and searching for food leaves a forager exposed to predators, yet forgoing opportunities to feed can incur risks in terms of reduced survival and reduced fitness through starvation (Olsson and Molokwu, 2008). Information about the level of predation risk, and the availability of food at a patch, could vary in quality over space and time, therefore foragers often have to make decisions about the costs and benefits of feeding based on incomplete information (Brown et al., 1999). Fixed habitat characteristics, such as trees and vegetation structure, influence perceived predation risk and could have a greater influence on foraging behaviours than more direct cues of predation, such as predator presence or density, perhaps because these offer more stable, but not necessarily more reliable, predictors of risk (Orrock, 2004; Verdolin, 2006). Here we investigate whether urban habitat features and human presence differentially influence foraging decisions of grey squirrels (*Sciurus carolinensis*) living at varying levels of urbanisation, with the hope to understand how urban features might influence foraging behaviour and perceived risk in a successful urban dwelling mammal. We consider the influence of proximity to trees, a static cue of risk known to influence grey squirrel perception of risk ( Newman and Caraco, 1987; Newman *et al*., 1988; Leaver, Jayne and Lea, 2017), as well as proximity to urban features, including roads and footpaths, which may offer more dynamic cues of risk relating to human disturbance.

The relationship between foraging, habitat use, and fear can be conceptualised via the ‘landscape of fear’. This suggests that habitat use is based on perceived risk or fear of predation (Laundré, Hernández and Ripple, 2010). Therefore, the way in which an animal uses its habitat can reveal much about the problems it faces in its day-to-day activities. Fear is a foraging cost through increased investment of time dedicated to vigilance and avoiding predators, thus reducing time for feeding and finding food (Brown, 1999). To optimise gains from feeding at a patch, foragers should feed for as long as the energy obtained is greater than the costs of foraging (Charnov, 1976). However, foragers assess more than the energetic value of a food patch. They also need to balance this with the costs of predation and the costs of missed opportunities to engage in other activities, such as finding mates and defending territories (Brown, 1988). As the forager depletes a food patch, these costs of foraging will begin to outweigh the benefits . The point at which this occurs can be quantified using the giving-up densities (GUD), the amount of food left behind after a forager stops feeding at standardised feeding patches ( Brown, 1999; Brown, 1988). Developed by Brown (1988) the GUD method has been employed to quantify foraging under risk and the landscape of fear in a variety of species, across diverse habitats and contexts. For example, the thermoregulatory costs of foraging (Kilpatrick, 2003), competition and niche separation (Brown et al., 1997) and responses to novel predators (Kovacs *et al*., 2012). GUDs reflect a foragers quitting harvest rate: the point at which survival and fitness costs begin to outweigh the energy gains ( Brown, 1999). As such, this point reveals a number of factors about the foragers and the foraging patch, including: level of predation risk , current energy state of the forager, foragers current fitness value, and the relative energetic value of the food ( Brown, 1988). As a consequence, GUD is expected to be high (more food left behind ) where predation risk is high and/or energy state and fitness value of forager “assets” are high, because foragers with a positive energy budget should be less willing to take foraging risks potentially impacting future fitness (*sensu* the asset protection principle: (McNamara, 1990; Clark, 1994).

With foraging behaviour revealing much about the survival and fitness consequences of local habitat use, examining foraging decisions in urban wildlife can offer a means of understanding impacts of human-altered environments on distribution, abundance, and selection pressures on wildlife inhabiting these landscapes. Mapping the landscape of fear in urban wildlife can also aid in development of urban wildlife management practices, for example, by identifying methods of artificially increasing the perceived predation risk to control impacts of foragers in certain areas (Abu Baker, Emerson and Brown, 2015). It is now widely accepted that human activity and related disturbance can significantly alter the behaviour of other species, with this is being most apparent in urban habitats (Lowry, Lill and Wong, 2013; Sol, Lapiedra and González-Lagos, 2013; Bates *et al*., 2015; Johnson and Munshi-South, 2017). Much of this change in behaviour might reflect differences in the perceived risk humans represent for a particular species. Humans can represent both an indirect and direct predation risk, similar to other apex predators, with their presence generating a landscape of fear in similar ways (Ciuti *et al*., 2012; Clinchy *et al*., 2016). For example, human activity and can increase mortality rates directly through vehicle collisions, destruction of habitat, or management of species regarded as pests. There are also indirect mortality costs associated with avoiding humans , including the energetic costs of increased physiological stress and exposure to disturbance (Ditchkoff, Saalfeld and Gibson, 2006). Whilst some species, may perceive human disturbance similarly to predation, human activity may actually reduce mortality for some, through reducing predator numbers or activity , perhaps providing a ‘safer’ habitat compared to their natural range (Fischer, Cleeton and Timothy, 2012)

Whilst there have been limited studies employing GUD methodology to assess human induced landscapes of fear within urban habitats ((Bowers and Breland, 1996; Fardell *et al*., 2022), other methods have revealed that spatial and temporal habitat use are shaped by human presence cues. In response to experimental playbacks of human voices, both large and medium-sized carnivores have demonstrated physiological and behavioural fear responses and avoidance of areas near cues of human disturbance (Clinchy *et al*., 2016). A study by Suraci, *et al* (2019) manipulated cues of human presence via playbacks throughout two 1km areas within the Santa Cruz mountains, USA, whilst monitoring the movements of mountain lions (*Puma concolor*) , meanwhile using camera traps and supplementary food patches to monitor movements of medium and smaller sized mammals including skunks (*Mephitis mephitis*), bobcats (*Lynx rufus*), opossums (*Didelphis virginiana*), deer mice (*Peromyscus* spp.) and woodrats (*Neotoma fuscipes*). Whilst mountain lions and medium-sized carnivores showed avoidance of areas with human playbacks, foraging by small rodents increased in these areas. This implies that, for these large and medium sized carnivore species, humans create a landscape of fear, which in turn provided predation release for smaller prey species. Furthermore, this provides an example of how human disturbance can alter and shape habitat use in whole communities of species, much like apex predators (Suraci *et al*., 2019).

A number of landscape of fear studies, demonstrate that it is not only predator species that avoid human activity (Bonnot *et al*., 2013; Rösner *et al*., 2014). Human recreational activity was found to impact patch use and GUD in Nubian ibex, *Capra nubiana,* (Tadesse and Kotler, 2012), revealing that human disturbance can represent a significant foraging cost. Similar costs of human disturbance have been highlighted by studies investigating the behavioural impacts of anthropogenic noise, where increased noise disturbance can lead to increased vigilance behaviours (Klett-Mingo et al., 2016; Sarno et al., 2014), spatial and temporal avoidance of foraging during increases in human activity levels (Luo, Siemers and Koselj, 2015), and additional energetic costs to other behaviours , such as mate selection and territorial defence, via making behavioural adjustments to disturbance (Fuller, Warren and Gaston, 2007; Barber, Crooks and Fristrup, 2010; Herborn *et al*., 2014; Morris-Drake *et al*., 2017; Petrelli *et al*., 2017).

To further contribute to the understanding of how urban living wildlife might change behaviours to minimise costs of human disturbance, here we investigate the patch choices in free ranging populations of eastern grey squirrels living at sites of varying urbanisation. A common invasive species in the UK, grey squirrels are abundant across landscape types and live at high density in many urban parks (Parker and Nilon, 2008; Bonnington, Gaston and Evans, 2013). Behavioural differences between urban and rural squirrels have become apparent within its native range, with urban squirrels showing reduced flight initiation distances towards human (Dill, 1989; Bateman and Fleming, 2014), living at higher population density (Sarno et al., 2014) and increased levels of intraspecific aggression (Parker and Nilon, 2008). Their success in urban habitats is thought to be due to their behavioural flexibility enabling them to adjust rapidly to novel problems (Chow, Clayton and Steele, 2021) and novel food resources, such as bird feeders and waste resources (Bonnington, Gaston and Evans, 2014; Kays and Parsons, 2014). This is likely to buffer them from the seasonal fluctuations in food supply usually experienced by woodland populations (Gurnell, 1996). Grey squirrels have been widely used in studies of foraging ecology and giving-up density (for examples see: Lima et al., 1985; Makowska & Kramer, 2007; Newman & Caraco, 1987; Wauters et al., 2000), so much is known already about their habitat preferences, however, research into their habitat use in urban environments appears to be limited, particularly in the UK, where their status as an invasive species means that further understanding of the factors influencing their distribution and abundance is important for wildlife management (Bonnington, Gaston and Evans, 2013, 2014).

In their native range , eastern grey squirrels inhabit densely forested habitat, preferring to feed and forage close to dense vegetation cover where they are protected from aerial predators ( Newman and Caraco, 1987). They are usually associated with habitats containing masting tree species, where their over-winter survival is impacted by the abundance of cashable food sources available in autumn (Fox, 1982). They coexist with other species of tree squirrel (e.g. fox squirrels *Sciurus niger* ) particularly in large urbanised parkland where a combination of anthropogenic food sources and mature masting tree species are thought to support high populations of squirrels (Van Der Merwe, Brown and Jackson, 2005)

In a study of urban squirrel foraging, Bowers and Breland (1996) used citizen science to measure the summer GUDs of grey squirrels at 78 artificial food patches placed individually at different points across an urbanisation gradient in Virginia, USA. They found a higher rate of locating artificial food patches and lower giving-up densities near areas of human habitation, suggesting that food patches near areas of human activity are readily exploited by squirrels, perhaps offering more abundant feeding opportunities and/or protection from natural predators (Bowers and Breland, 1996). Here, as well as measuring GUDs along a UK urban-rural gradient, we investigate how different habitat features associated with urbanisation influence this. Given previous findings from GUD studies in grey squirrels, perceived foraging risk will likely increase with distance from trees and vegetation cover, therefore we provide foragers with ‘safe’ (close to trees) and ‘risky’ (further from cover) pairs of foraging patches at each site along the gradient. This design assays the local perceived predation risk with greater differences in GUDs expected under higher levels of perceived predation risk at a site, as the foraging value of being close to cover will be higher (safer patches will be exploited more intensively) under riskier conditions. Such a paired design controls for variation in local food availability and other ecological influences on the costs (including lost opportunity costs) of foraging at any given site (Brown 1988). However, whether a feature represents a stable or dynamic cue of disturbance may influence differences in safe vs risky GUDs further. We measure distance of each feeding patch pair from roads, footpaths, and buildings, with roads and footpaths thought to present more dynamic cues of human disturbance and buildings more static cues. In addition, we quantified local noise levels to examine if this mediates the relationship between GUD and proximity to urban features. We considered both average noise levels and noise variability as variable noise disturbances are predicted to disrupt foraging, for example through startle responses, or distraction (Chan *et al*., 2010; Gil, Klett-mingo and Pav, 2016; Evans, Dall and Kight, 2018) and animals may be less likely to habituate to stimuli that are variable or inconsistent (Rankin *et al*., 2009).

Therefore, we predicted that: 1) GUD differences between ‘safe’ and ‘risky’ patches would be greater at less urbanised sites, 2) proximity to features representing dynamic cues of disturbance (roads and footpaths) would increase the difference in GUDs within pairs.

## Materials and methods

### Study sites

The study was conducted during winter (December 2014-Febuary 2015 for four of the sites, and December 2015 – February 2016 for the remaining Arboretum and Pepperboxes Woodland sites) at six sites across Buckinghamshire, UK, varying in levels of urbanisation (Table 1). Data were collected during winter as natural sources of food are scarce at this time, therefore squirrels spend more time foraging at ground level (Parker, Gonzales and Nilon, 2014).Daily temperatures during data collection ranged between -4°C to 9°C. Attempts were made to ensure temperature and weather conditions were similar on the days prior and during GUD data collection in order to keep temperature related energetic costs of foraging as similar as possible across sites. Sites were all located within a 15 km radius of central High Wycombe, UK. Locations were initially selected using satellite images from Google Earth (version 7.1) based on proximity to urban built structures and containing suitable habitat for squirrels (all site contained some tree canopy). After confirming that squirrels were present and observable at selected sites, two ‘urban’ sites, two ‘suburban’ and two woodland sites were chosen (Further information on sites contained in supplementary information, S.1). . All sites contained potential predator species, including red fox (*Vulpes vulpes)*, red kites (*Milvus milvus*), and buzzards (*Buteo buteo*) were also reported at the Pepperboxes woodland site. Although abundance of domestic dogs (*Canis familiaris)* and cats (*Felis catus)* were not measured in this study, it is likely that they are found at higher density in urban and suburban areas (Baker *et al*., 2005; Weston *et al*., 2014).Sites were at least 5 km from each other, therefore it is likely that these represent independent populations. Daily home range sizes for grey squirrels are reported to cover 1.20 acres (∼4.85 km^2^) with a maximum linear distance of 136.7 m (Doebel and McGinnes, 1974), however, there have been cases of dispersal distances up to 64 km (Rushton *et al*., 1997), as a consequence we cannot be certain there were no study squirrels that did not visit another site that was included in this study, although it is unlikely.

**Table 1.** Characteristics of study sites.

| Site | Mean<br>building<br>density | No.<br>squares<br>>50%<br>build | Mean<br>vegetation<br>density | No.<br>squares<br>>50%<br>veg | No.<br>road<br>squares | Urban<br>index<br>(PC1) | Squirrel<br>population<br>estimate<br>(individuals/acre) | Urban<br>Category |
| --- | --- | --- | --- | --- | --- | --- | --- | --- |
| Naphill Forest | 0.06 | 0 | 1.99 | 98 | 4 | -48.81 | 0.19 | Rural |
| Pepperboxes<br>Wood | 0.25 | 3 | 2.02 | 100 | 28 | -36.19 | 0.8 | Rural |
| Kingshill<br>Arboretum | 0.75 | 16 | 0.96 | 96 | 34 | -23.69 | 0.85 | Rural |
| Wycombe Park | 0.48 | 17 | 1.36 | 44 | 24 | 7.28 | 4.14 | Urban |
| Missenden<br>Gardens | 0.79 | 20 | 1.4 | 40 | 27 | 13.11 | 0.85 | Urban |
| Wycombe<br>Museum | 1.7 | 72 | 0.73 | 16 | 89 | 88.28 | 1.07 | Urban |

Land cover scores for 100 squares (100mX100m) covering grid of 1-km<sup>2</sup> area around each site. PC1 values calculated from principle component analysis on the five variables for each site using methodology described in (Czúni, Lipovits and Seress, 2012). Larger scores indicate higher building and road cover and less vegetation cover.

To enable sites to be broadly categorised and compared in terms of level of urbanisation, three main features of land cover type: roads, buildings and vegetation cover were used to provide an index of urbanisation for each site. Using digital images in Google Earth Pro, sites were divided into 10X10 grid (100 squares of 100m X 100m), covering a total area of 1-km^2^ around (including and surrounding) each site. Each square was scored according to building cover (0 if ratio of buildings is 0%; 1, if < 50%; 2, if >50%), vegetation cover (0, if 0%; 1, if < 50%; 2, if >50%) and if solid road was present (0, if absent; 1, if present). An urbanisation index score for each site was then created from the first principle component score calculated using the summary statistics of the 100 squares at each site (Liker *et al*., 2008; Czúni, Lipovits and Seress, 2012). PCA was carried out using the *prcomp* function in R version 3.5.2.

To aid in our interpretation of GUD, estimates of squirrel population density surrounding each site were carried out to provide a proxy measure for potential food competition levels at each site. Whilst population density is not often included in GUD studies (Carthey and Banks, 2015), the density of conspecifics is likely to increase competition for food resources, and may generate lower GUDs. Grey squirrels were observed regularly at all six sites and their local abundance in the South East of England has been noted since at least 1916 (Middleton, 1930). Fixed-duration point counts were used to provide an estimate of squirrel abundance as this provided a way to calculate an index of population density in fragmented urban sites where line transects are not always possible (Parker and Nilon, 2008; Bonnington, Gaston and Evans, 2013). Following similar methodology to that described (Flyger, 1959), 15min point counts were made from 10 observation points covering the area within and surrounding the GUD grid, with each of these points being a minimum of 136m apart to account for home range sizes (Doebel and McGinnes, 1974) reducing the likelihood of repeated counts of the same individuals. Individual points were selected via an initial walking transect stopping every ∼150m, where the nearest suitable observation point with a 360-degree visual field was selected. These transects usually followed existing footpaths paths and trails used by walkers. Counts at each point were conducted at least four times at each site prior to collection of GUD data, and were carried out on dry calm days (<4 Beaufort scale) within five hours of sunrise when squirrel activity is usually highest (Parker and Nilon, 2008). (Further information contained in supplementary information, S.2)

Squirrel densities ranged from 0.19 – 4.14 individuals per acre, with largest estimates for population densities occurring at the urban park site (Table 1). In their native range densities of grey squirrel populations typically range from 0.3 – 0.8 individuals/ha (0.75 – 2 individuals/acre) (Williamson, 1983). With UK urban populations found to range from 0.46 – 8.29 individuals/ha (1.15 – 20.7 per acre) (Bonnington, Gaston and Evans, 2013).

### Measuring giving-up densities (GUDs)

At each site, an area of 100m^2^ was selected based on observations of squirrel feeding activity, as well as accessibility and landowner permissions. Within this area, eighteen food patches were placed in pairs across a 3X3 grid, creating nine feeding locations spaced a minimum of 30m apart (N = 108 patches across all sites). At each of these locations food trays were placed in pairs with one tray at the base of the closest mature tree with a minimum diameter at chest height of 45cm (Parker and Nilon, 2008), and the other tray placed 3-7 meters away in open habitat, at least 3 meters from dense ground vegetation cover. For seven days squirrels were habituated to feeding from the trays before data collection begun. This was done to reduce potential impacts of differences in neophobic responses, as we might expect urban squirrels to be more familiar with obtaining food from novel anthropogenic food sources than forest dwelling squirrels. Feeding signs, such as footprints, peanut skins and sunflower shells (from bait provided), were used to assess whether squirrels were encountering the trays. If trays showed no feeding signs for more than 3 days in a row, they were moved to a new location in the surrounding area.

Giving-up density was measured using the weight of food left behind at each artificial food patch. These feeding patches consisted of green plastic trays (38cm L x 24cm W x 5cm high) each containing 15 grams of granulated peanuts buried in 2.5 litres of sand. Granulated peanuts were used because squirrels consumed these at patches, rather than cashing single nuts. A grid constructed of plastic mesh with 5cmX5cm squares was placed on the top of each tray to further increase diminishing returns as squirrels foraged.. To reduce foraging by birds trays were also covered with a cage of wire mesh allowing squirrels to enter and exit on two sides. Patches were opened at sunrise and closed at sunset (approx. 0800 – 1600) during this study, therefore the weight of peanuts left in the tray reflected 8 hours of foraging. Trays were sieved to separate the peanuts from the sand substrate, and the weight of peanuts remaining was recorded as the giving-up densities.

### Measures of proximity to urban features and anthropogenic disturbance

GPS coordinates for each tray pair were entered into EDINA DigiMap (EDINA DigiMap, Edinburgh, UK) and measurements from the mid-point of each pair of trays to the nearest building, road, and footpath were taken using the distance ‘*Roam*’ measurement function.

At each pair of food patches, noise levels were monitored from a point equidistant between each tray. Noise levels were sampled for ∼15 seconds (with a dB reading noted approximately every second) at three time points each day (morning ∼0800 – 0930, afternoon ∼1200 – 1400, evening ∼1600-1800) over three days using a hand-held Benetech digital sound level meter (A-weighted, range 30 – 130 dB). Data points were noted by a single observer with the sound level meter held at a hight of ∼1-1.20 m from ground level.This resulted in 135 sampling points for each tray pair. The mean and variance in noise levels were calculated for each pair in an attempt to capture differences between patches in both average levels of noise as well as exposure to intermittent noise disturbances.

### Ethical note

Research followed and adhered to the ASAB guidelines for treatment of animals in behavioural research (ASAB, 2020). The project was reviewed and approved by the University of Exeter Biosciences Review Panel. Feeding trays were cleaned at the end of each day and replaced with fresh substrate and food to reduce potential for exposure to disease or parasites. Pre-baiting and feeding lasted for no more than 10 days, unlikely to cause significant changes to squirrel population or reliance on this as a supplementary food source.

## Analysis & Results

Squirrels left higher GUDs at food patches further from cover (F1) 28.68, p<.001) (Table 2); with urban sites showing lower GUDs compared to rural sites overall (F(1) 31.96, p<.001).

**Table 2:** Mean GUDs at each site. Sites ordered from least to most urban.

| Site Name | Site Category | Mean (SD) GUD Safe patches (g) | Mean (SD) GUD risky patches (g) |
| --- | --- | --- | --- |
| Naphill Forest | Rural | 7.346 (3.141) | 9.593 (3.744) |
| Pepperboxes Wood | Rural | 5.457 (2.964) | 11.753 (1.719) |
| Arboretum | Rural | 6.756 (2.639) | 8.647 (3.738) |
| Park | Urban | 2.252 (0.870) | 5.771 (1.924) |
| Gardens | Urban | 3.663 (1.688) | 8.851 (4.304) |
| Museum | Urban | 4.766 (4.009) | 5.956 (5.159) |

Generalised linear mixed models with gaussian error were used to test for the effects of proximity of urban features and noise levels on the differences in GUD between open and cover patches in each pair. GUD difference (ΔGUD) was calculated by subtracting the GUD at the patch further from cover (open) from the GUD at the patch close to the base of a tree (Cover). Higher ΔGUD scores indicate that cover patches were exploited more intensively than open patches thus providing an index of the relative predation costs of foraging for squirrels at each pair of patches. Prior to analysis variables were checked for normality. Three data points in GUD that were found to fall above or below the inter-quartile range were removed from the data set, these were identified using the R function *box.plot.stats().* Proximity to urban features, mean and variability in noise levels were rescaled and mean centred. Urban feature variables (distance from roads, footpaths, and buildings) were examined for collinearity using Pearson correlations. Roads and buildings were found to be highly correlated (r = .95), therefore distance from buildings was removed from the model. Distance from roads was retained, as it might be considered the more direct index of disturbance risk.

The maximal model contained continuous predictor variables: distance from roads distance from footpaths, noise variance, (mean) noise level and their two-way interactions. Site was included as a random effect. All analyses carried out using R version 3.5.2. Linear mixed models were carried out using the *lmer* in the package ‘*lme4’* (Bates *et al*., 2015), model selection were carried out using backwards stepwise regression (Fox and Weisber, 2019). Terms were excluded based on whether their exclusion increased each model Akaikes Information Criterion (AIC). Confidence intervals associated with each fixed effect were calculated using the *confint* function.

The minimal model (Table 3) retained the fixed factors: distance from roads, noise level and noise variance. There were significant interactions between distance from roads and noise levels , t(43) -2.280, p = .027, and between distance from roads and noise variability, t(43) 2.86, p = .006, on ΔGUD. Foragers feeding closer to roads left smaller differences in GUD between safe and risky patches (i.e. they perceived lower levels of predation risk) when noise levels were high (Figure 2), although where variability in noise levels increased, so did ΔGUD (Figure 3) (suggesting they perceived higher levels of predation risk when noise levels are variable). No significant main effects of these variables were evident (Table 3).

**Figure 2:**
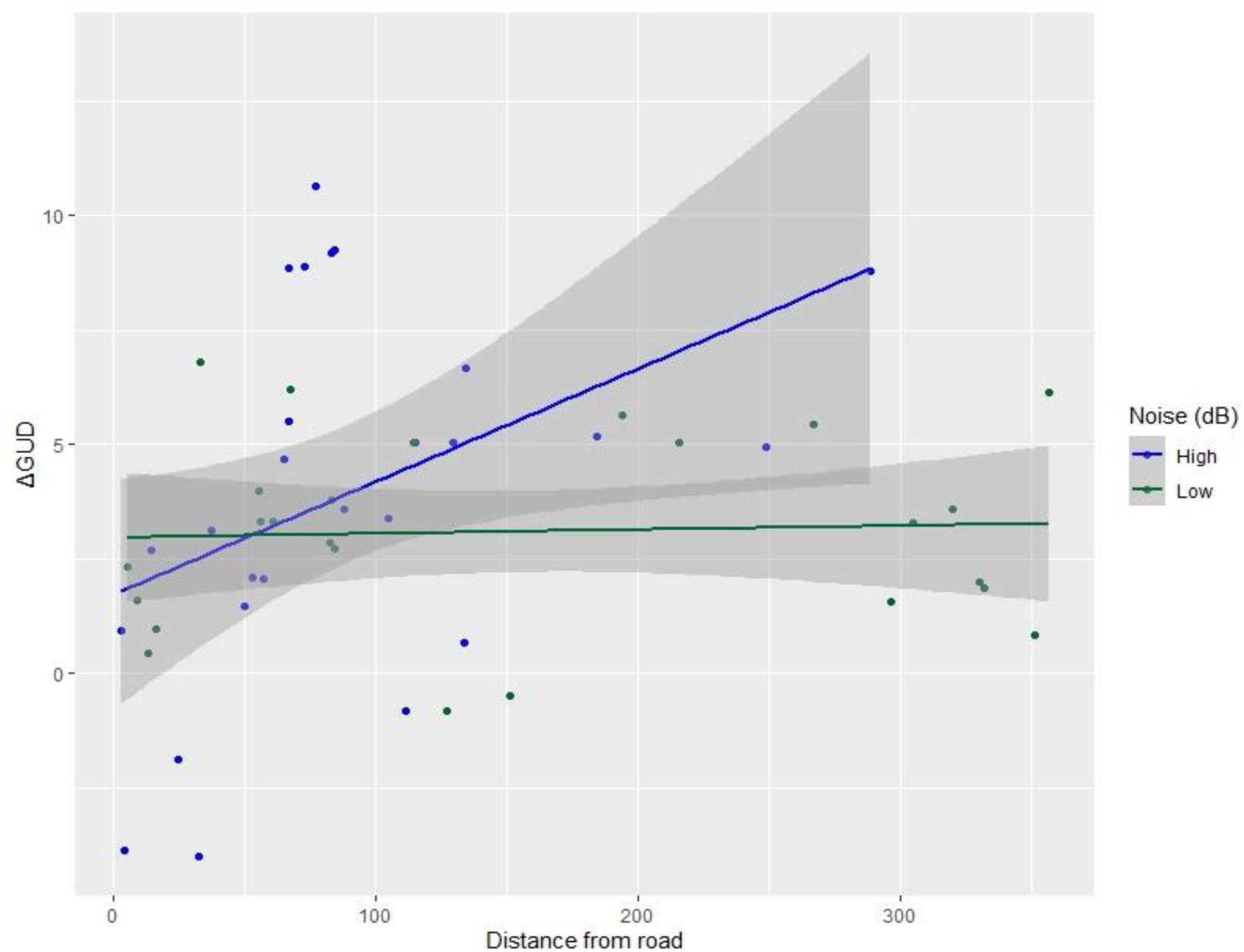
Interaction between distance from roads and noise levels on ΔGUD *Noise level was split into categories based on a median split.* Under high noise levels, ΔGUD (open/risky patches – cover/safe patches) increases with distance from roads, suggesting that under high noise conditions, squirrels preferred to forage at patches closer to trees (cover), with this preference increasing as distance from roads increases, suggesting foraging is perceived as riskier under high noise levels with increasing distance from roads.

**Figure 3:**
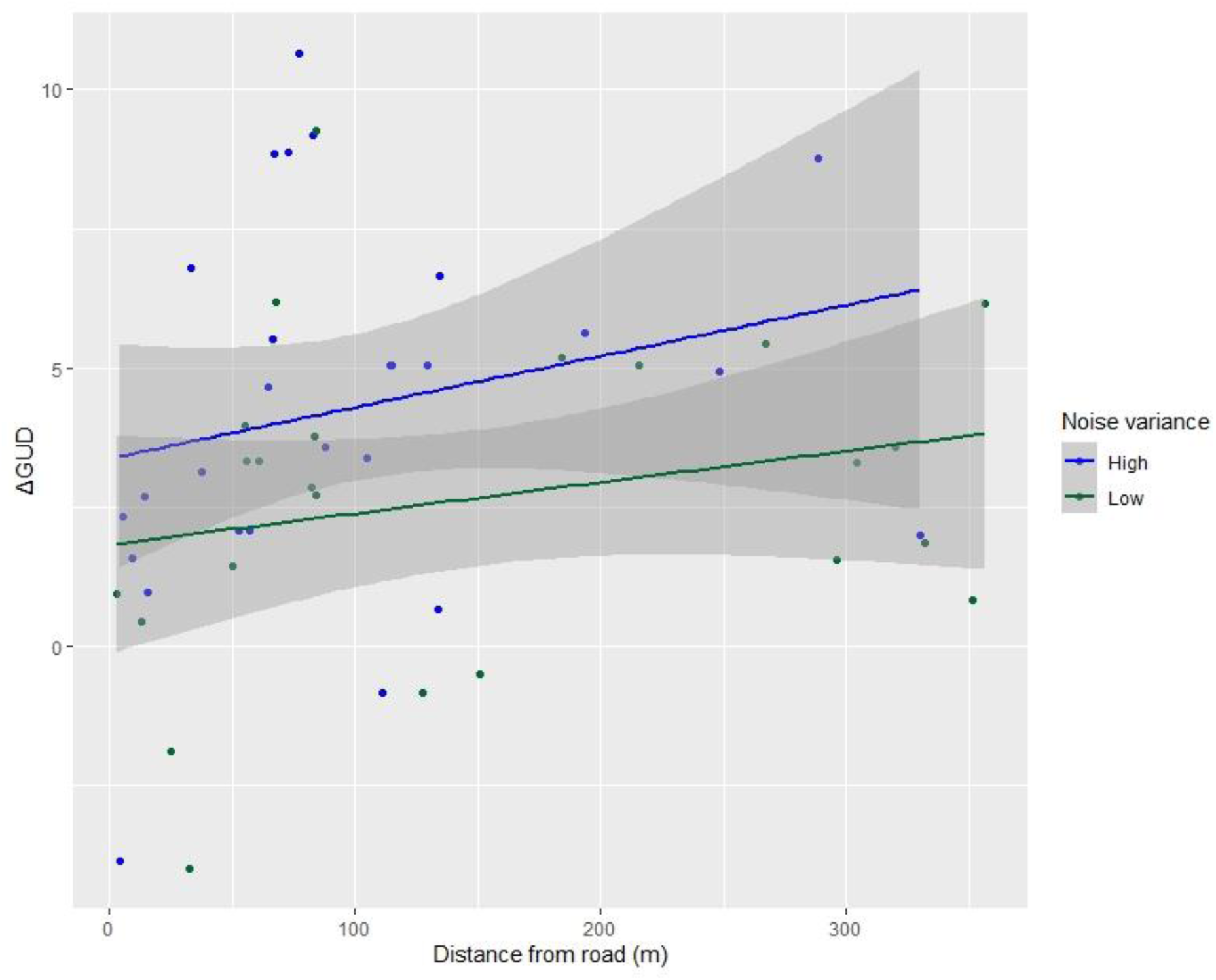
Interaction between distance from roads and noise variability on ΔGUD *Noise variance was split into categories based on a median split.* Under both high and low levels of noise variance percieved risk increases with a greater distance from roads. However, the effect of proximity to road on ΔGUD appears enhanced under higher noise variance conditions.

**Table 3:** Summary of fixed factors included in LMM full and minimal adequate model influencing GUD difference between safe and risky patches. Site included as random factor.

| Full Model (AIC 259.29) |  |  |  |  |  | Minimal Model (AIC 254.40) |  |  |  |  |
| --- | --- | --- | --- | --- | --- | --- | --- | --- | --- | --- |
| Variable | Estimate (±SE) | df | t | p | CI | Estimate (±SE) | df | t | p | CI |
| (Intercept) | 4.506<br>(0.707) | 5.493 | 6.375 | >0.001 | (3.308 - 5.762) | 4.4797<br>(0.689) | 5.722 | 6.494 | >0.0001 | (3.273-5.699) |
| Road distance | 1.269<br>(0.820) | 36.210 | 1.547 | 0.130 | (-0.077 -3.070) | 1.219<br>(0.693) | 38.351 | 1.759 | 0.086 | (0.0328 - 2.738) |
| Footpath distance | 0.127 (0.454) | 40.6 | 0.279 | 0.781 | (-0.829-0.895) |  |  |  |  |  |
| Noise Variance | 0.587 (0.570) | 41.731 | 1.029 | 0.309 | (-0.663 - 1.527) | 0.539 (0.522) | 44.798 | 1.031 | 0.308 | (-0.625 - 1.448) |
| Noise level | 1.497(1.169) | 39.390 | 1.280 | 0.207 | (-0.439 - 3.924) | 1.565 (1.071) | 43.856 | 1.462 | 0.151 | (-0.318 - 3.829) |
| Noise var X Road distance | -1.82 - 0.963) | 41.3 | -1.897 | 0.064 | (-3.812 - -0.196) | -1.957 (0.858) | 43.843 | -2.280 | **0.027** | (-3.732 - 0.409) |
| Noise var X footpath distance | (-0.326 - 0.492) | 39.502 | -0.669 | 0.507 | (-1.272 - 0.565) |  |  |  |  |  |
| Noise level X Road distance | (3.307 - 1.547) | 35.630 | 2.137 | **0.039** | (0.766 - 6.759) | 3.312(1.157) | 43.74 | 2.862 | **0.0064** |  |
| Noise level X footpath distance | (-0.005 - 0.708) | 41.946 | -0.008 | 0.993 | (-1.517 - 1.170) |  |  |  |  |  |

## Discussion

In this study we found that invasive grey squirrels across all sites showed predation risk sensitive patch use, on average leaving lower GUDs closer to trees than in open (risky) habitat. This reflects the microhabitat preferences found in GUD studies conducted within their native range (Lima, Valone and Caraco, 1985) where foraging near the base of large trees offers both an escape route from terrestrial predators and canopy protection from predatory birds ( Newman and Caraco, 1987). We assayed GUD differences within matched pairs of artificial food patches (near trees vs open) to assess if anthropogenic habitat features influence the differential exploitation of safe versus risky patches and thereby the landscape of fear in our study population. We found that the difference between the exploitation levels of safe and risky patches decreased closer to roads, suggesting that the squirrels in our study perceived roads as ameliorating predation risk. Indeed, this relationship between proximity to roads and perceived risk was found to be significantly impacted by noise. The mitigation of perceived predation risk by proximity to roads was particularly pronounced when noise levels were high (Figure 2) suggesting that the squirrels in our study perceived themselves as safest from predators where disturbance by human activity is likely to be particularly high. However, the squirrels in our study also increased their differential exploitation of safe patches under more variable noise levels, increasing their foraging at safe patches (reducing foraging in risky patches) under higher levels of noise variance. Although under both low and high noise variance, differential GUD decreased with proximity to roads, the ameliorating effect of roads appeared slightly enhanced under higher noise variance conditions (Figure 3). Thus some human activities that generate inconsistent noise may also act to increase perceived predation risk in invasive grey squirrels, perhaps by impacting reliability of cues of predator activity (Warren *et al*., 2006; Gill *et al*., 2015). No significant effects of distance to footpaths and buildings were found, although broadly, squirrels feeding at more urban sites (those containing more buildings and roads) left lower GUDs suggesting that squirrels at urban sites perceive predation risk as lower.

Whilst these results suggest grey squirrels may perceive sites close to roads as safer, this may not necessarily imply that living close to roads and areas of human disturbance is risk free. Roads are generally associated with the mortality risk posed by collisions with vehicles (Ware *et al*., 2015). Roadkill surveys in the UK and USA suggest that roads do present a significant mortality risk for grey squirrels, and that this may vary spatially and temporally (Smith-Patten and Patten, 2008; Raymond *et al*., 2021). For example, in the UK, grey squirrel roadkill reports show peaks occurring in summer and autumn, times when dispersal and foraging activity are expected to be high for this species (Raymond *et al*., 2021). Some animals living close to roads may be able to adjust behaviour in response to perceived risk of vehicles, for example, by adjusting flight initiation distances relative to vehicle speed (Legagneux and Ducatez, 2013; Lunn *et al*., 2022). For prey species, the noise produced by traffic may mask detectability of predators, including heterospecific or conspecifics alarm calls, leading some animals living close to roads becoming less cautious during foraging (Giordano, Hunninck and Sheriff, 2022) whilst others to increase vigilance (Merrall and Evans, 2020) . Road noise could also impact predator behaviour, through mortality risk and reducing the ability to detect prey, which in turn may lead prey to be less cautious (Chan *et al*., 2010; Giordano, Hunninck and Sheriff, 2022). Studies using road noise playbacks suggest that road noise reduces foraging efficiency, as animals invest in increased vigilance(McClure *et al*., 2013; Ware *et al*., 2015). However, it is possible some species will be more likely to habituate or cope with this background noise than others(Uchida and Blumstein, 2021), and it is likely these impacts will be further influenced by how food resources and predator activity may be affected by levels of urbanisation and type of human activity in certain areas.

Urban habitats may contain altered and novel sources of mortality , although actual predation rates in urban environments appear to be lower (Fischer, Cleeton and Timothy, 2012). Sites near human settlement and disturbance may offer a refuge from natural predators for some species, whilst encounters with humans are likely to be non-fatal in general, and this may explain why, in this study, squirrels were able to forage for longer. Previous studies of urban grey squirrels have found that urbanised squirrels become less wary of human activity showing reduced flight initiation distances as humans approach (McCleery, 2009; Parker & Nilon, 2008), although the level of predictability, type and frequency of human disturbance appears to cause variability in the vigilance behaviours of grey squirrels, suggesting that not all forms of human disturbance are perceived equally in terms of predation risk (Sarno et al., 2014). Furthermore, in urban settings, the ‘safe’ patches, close to vegetation and trees used in this study may not necessarily represent a safer foraging option, as it would in grey squirrels’ native range. Urban gardens may contain higher densities of domestic pets, altering predation risk. For example, domestic cats are ambush predators (Baker *et al*., 2005), therefore foraging in open habitat may be safer for prey species where cats are present. Further research could benefit from collecting detailed data on the abundance and distribution of predators at each site to aid interpretation of results.

Lower GUDs observed in urban sites could also reveal differences between sites in terms of resource quality and missed opportunity costs. Foraging models based on missed opportunity costs predict that GUDs should be higher in environments with high food availability and short travel distances between patches (Olsson and Molokwu, 2008). This may provide an explanation for the higher GUD observed in forested sites, where an abundance of masting trees, such as beech and oak, may provide plentiful alternative foraging opportunities at the time data was collected. In contrast, urban sites may require further, energetically costly, travel distances between aggregated food patches (Hinsley *et al*., 2008), resulting in increased foraging intensities (lower GUDs) due to the increased in value of any patches that are encountered. In a habitat containing high food availability, the differences in GUDs between safe and risky patches are predicted be greater as foragers are less likely to take risks (Olsson and Molokwu, 2008). Conversely, where food availability is limited or patchy, and/or where the foragers are in a low energy state, food becomes a higher value resource and foragers more may be more willing to take greater risks to acquire it (Namara, 1990). Therefore, the reduced differences in GUDs between safe and risky patches close to urban features may reveal that, in urban habitats, squirrel foraging behaviour shifts from being predator-limited, to being food-limited (Bowers and Breland, 1996).

Based on our squirrel population estimates, we found that sites scoring higher in urbanisation levels contained higher squirrel densities. It is possible that higher population densities at these sites lead to lower GUDs perhaps due to increased competition for food resources, or due to the potential benefits of feeding close to other foragers. Grey squirrels are not regarded as particularly social or territorial in comparison to other sciurids (Koprowski, 1996), however, they demonstrate dominance hierarches in the context of resource access (Allen and Aspey, 1986), and forage in the presence of conspecifics (Hopewell and Leaver, 2008). The increased presence of conspecifics in habitats of higher population density may allow squirrels to use social information, rather than habitat features, to make assessments about risk (Lima, 1995). Foraging grey squirrels are known to respond to feeding, vigilance, and alarm call behaviour in both conspecific and heterospecific foragers (Partan *et al*., 2010; Getschow *et al*., 2013; Jayne, Lea and Leaver, 2015). In a study of urban grey squirrel responses to conspecific alarm calls, it was found that urban squirrels were more reactive to alarm signalling than less urbanised squirrels, and were found to attend most strongly to visual alarm signalling, such as tail flagging, perhaps due to the impact of anthropogenic noise disturbance on attention to auditory signals (Partan, Larco and Owens, 2009). As such, foraging within a group may reduce perceived predation risk for grey squirrels through the benefit of the information provided by ‘many eyes’ (Lima, 1995). This could allow individuals to reduce vigilance behaviours and invest more time in foraging and feeding, resulting in lower GUDs. Furthermore, it may allow foragers to exploit ‘risker’ patches more readily if conspecifics are present. Although the effects of social foraging on GUD has not been as widely studied , in a study of the GUD of black rats (*Rattus rattus*) and bush rats (*Rattus fuscipes*), it was found that solo foragers left higher GUD than those foraging in a group, suggesting that social context may be an important factor to consider in GUD studies (Carthey and Banks, 2015).

Whilst reducing perceived predation risk, the presence of other foragers can represent additional foraging costs including patch defence and the risk of aggressive interactions. It is possible that through exploitative competition, individuals in urban sites may be driven to forage more rapidly to obtain a greater share of a high value resource. Furthermore, it may be that the individuals more willing to risk antagonistic interactions with conspecifics are those willing to forage at patches close to areas anthropogenic disturbance. Further work would benefit from taking an individual based approach to find which individuals are foraging at these patches. For example, it might be that bolder individuals, less sensitive to human disturbance, dominate these food patches. We may also find that it is individuals in a low energy state due to higher competition for resources in urban habitats (Shochat and State, 2007), that are more likely to exploit patches near sites of human disturbance and engage in riskier foraging. Data collected in this study took place just within typical breeding season: January-June for UK grey squirrels ( (Middleton, 1930), therefore the value of mating opportunities is likely to be high during some of the time data was collected. According to the asset protection principle (Clark, 1994), the value of safety becomes higher in individuals with high reproductive potential. Squirrels living in sites further from human disturbance could be in a higher energy state, or better body condition, so may be less willing to take foraging risks. However, further studies are needed to assess differences in body condition, reproductive success, and survival rates of grey squirrels across the urbanisation gradient in order to put these findings in this context.

Overall, the results in this study suggest that foraging costs and perception of risk in foragers are altered by proximity to roads and variability in noise disturbance: key urban features related to human activity. This can be interpreted as there being lower predation risk close to roads, or that the marginal value of food is higher for foragers at these sites. It is likely that the differences in GUD across sites in this study reveal a variation in the balance between predation risk, food value and missed opportunity costs across different levels of urbanisation. Future research into foraging and patch use across urbanisation levels should examine the variations in food supply, body condition, reproductive and survival differences in squirrels across disturbance gradients, areas that appear understudied in UK populations of grey squirrels, as this could aid in a better understanding and interpretation of differences in behaviour, distribution and abundance of squirrel populations living at varying levels of human disturbance.

