## Supplemental Information for "The double-edged sword of urban life: Invasive grey squirrels perceive foraging as safer close to roads under noisier conditions, yet riskier when noise is inconsistent"

**S.1 Description of Study sites.**

Urban sites were located within the town of High Wycombe, UK (Wycombe Museum and Wycombe School Park sites) were considered urban centre sites. Both sites contained mature beech trees (*Fagus sylvatica)* as well as limes (*Tilia europaea)* and yew (*Taxus baccata)*. The two 'suburban' sites consisted of one site comprised of five adjacent private gardens (labelled Missenden Gardens, Table 1), and an Arboretum in Little Kingshill, Buckinghamshire, UK. Forested sites consisted of a 33.11 acre (Pepperboxes wood) and a 155 acre site of ancient woodlands (Naphill Forest). Dominant tree species include mature European Beech (*Fagus sylvatica)*, Ash (*Fraxinus excelsior* ) and Oaks (*Quercus robur)*.

Bird feeders were observed at the garden site, and members of the public were often observed feeding squirrels at the urban park and museum site.

| **Site name** | **Latitude-longitude** | **Categorisation** |
|---|---|---|
| Wycombe Museum (red) | 51°37'52"N 0°44'54"W | Urban |
| Wycombe Park (red) | 51°37'22"N 0°44'33"W | Urban |
| Missenden Gardens (blue) | 51°41'20"N 0°42'30"W | Suburban |
| Arboretum (blue) | 51°41'08"N 0°41'51"W | Suburban |
| Pepperboxes wood (green) | 51°42'32"N 0°44'53"W | Woodland |
| Naphill Forest (green) | 51°40'05"N 0°47'28"W | Woodland |

*Image 1: Sites. Image via Google Earth*

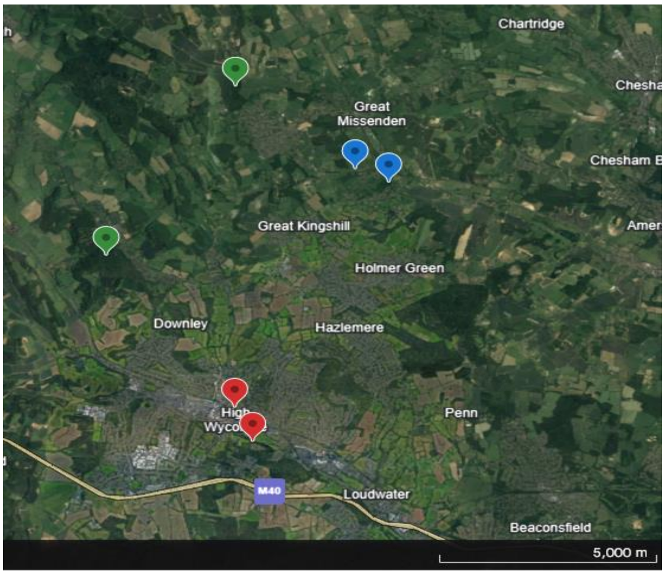


**S.2 Squirrel point-count calculation.**

Squirrel counts begun as soon as the observer approached the observation point. Using a range finder (Viking monocular laser range finder, Viking Optical, Suffolk, UK) the radial distance of each squirrel from the observer was estimated. Using the following equation, an index of relative population density (N) was calculated from the total area of the site covered (A), total number of squirrels counted (Z), the number of 15min observation sessions (n) and the average radial distance to squirrels counted ( r) (Flyger, 1959; Parker and Nilon, 2008):

$$N = \frac{AZ}{n\pi r^2}$$

Whilst our chosen method attempts to minimise the repeated sampling of the same individuals, this may have occurred. This method of estimating population density may also be subject to inaccuracy as it relies on one observer's ability to spot and count squirrels. Future research could benefit from more accurate methods for including population size, such as trap-mark-recapture procedures, however, this method does allow us to gain an approximation of population size that can aid in interpreting GUD outcomes across sites.

**S.3 GUD feeding tray set up.**

Feeding trays were trialled prior to the study, to ensure they contained suitable ratios of substrate to food. Using a camera trap on a trial GUD set up revealed that corvids were also feeding from the trays. Wire mesh was then added to prevent corvids.

*Image 2: GUD patch*

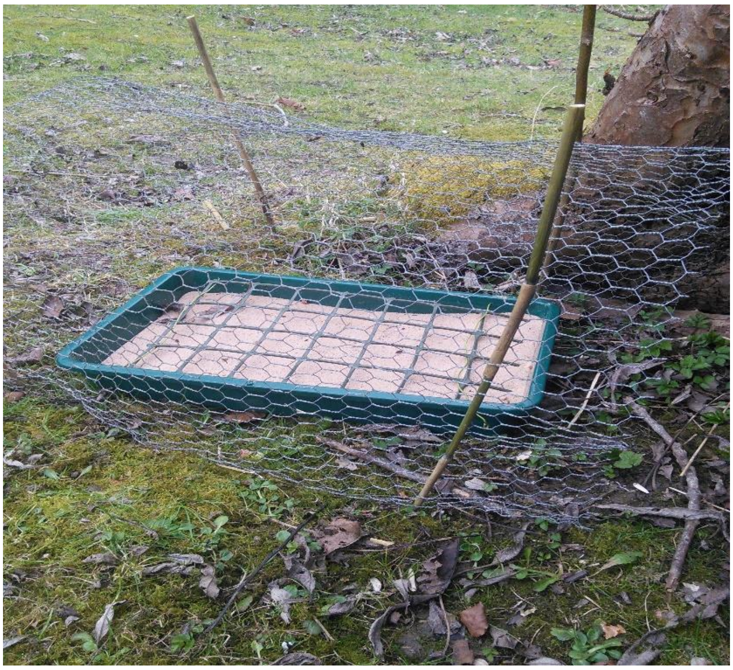
